# ChipFilter: Microfluidic Based Comprehensive Sample Preparation Methodology for Metaproteomics

**DOI:** 10.1101/2023.01.18.524548

**Authors:** Ranjith Kumar Ravi Kumar, Massamba Mbacke Ndiaye, Iman Haddad, Joelle Vinh, Yann Verdier

## Abstract

Metaproteomic approach is an attractive way to describe a microbiome at the functional level, allowing the identification and quantification of proteins across a broad dynamic range as well as detection of post-translational modifications. However, it remains relatively underutilized, mainly due to technical challenges that should be addressed, including the complexity in extracting proteins from heterogenous microbial communities.

Here, we show that a ChipFilter microfluidic device coupled to LC-MS/MS can successfully be used for identification of microbial proteins. Using cultures of *E. coli, B. subtilis* and *S. cerevisiae*, we have shown that it is possible to directly lyse the cells and digest the proteins in the ChipFilter to allow higher number of proteins and peptides identification than standard protocols, even at low cell density. The peptides produced are overall longer after ChipFilter digestion but show no change in their degree of hydrophobicity. Analysis of a more complex mixture of 17 species from the gut microbiome showed that the ChipFilter preparation was able to identify and estimate the amount of 16 of these species.

These results show that ChipFilter can be used for the proteomic study of microbiomes, in particular in the case of low volume or low cell density.

## INTRODUCTION

From the introduction of metaproteome and metaproteomics concepts (Rodriguez-Valera, 2004; Wilmes & Bond, 2004), studies have been done on various microbial communities. For examples, intestinal microecology, marine biology, soil biology, aerosols composition, and studies of food composition to explain food quality, safety and allergies (Yang et al., 2020). One of the major advantages of metaproteomics studies is that they provide functional information and reveal the microorganism functions and interactions at the protein level, complementary to metagenomic or metatranscriptomics data (Kleiner, 2019, Heintz-Buschart & Wilmes, 2018). Yet, metaproteomics remains relatively underutilized, mainly due to the challenges that remain in extracting proteins from heterogeneous microbial communities.

Extracting proteins from different microbial communities includes various challenges, without even mentioning sampling problems. While a universal extraction protocol providing good protein yields from a wide range of samples would be desirable, this objective does not seem achievable given the heterogeneity of matrices and microorganisms’ characteristics (Keiblinger et al., 2016). Commonly, protein extraction includes a direct cellular lysis step, which is attained via chemical lysis using detergents and stabilizing agents; physical lysis (heat, pressure or snap-freezing) or enzymatic lysis. The choice of the detergent is crucial and greatly impacts on the quality of the results (Glatter et al., 2015). Proteins can be purified using different methods, such as filter-based methods FASP (Wiśniewski et al., 2009), precipitation with acids like trichloroacetic acid (TCA) (Pérez-Rodriguez et al., 2020), separation on a polyacrylamide gel using electrophoretic mobility (Granvogl et al., 2007), or solid-phase separation (Hugues et al., 2014). Each method has advantages, like protein fractionation, recovery and washing off the detergent. Then, proteins are in most cases identified by mass spectrometry (MS) using a bottom-up approach, in which proteolytic peptides are analysed to generate protein inferences. The choice of the pre-analytical strategy must consider the heterogeneity of microbial cells, having varied cellular membranes that cannot be lysed by a universal method and have species specific challenges such as a high nucleic acids contents for bacteria or wall structure for fungi. Furthermore, combining several cell lysis procedures increases the risk of losing low-abundant proteins, experimental time and handling steps. Other challenges include automation, repeatability between biological replicates and interference of detergents with the subsequent purification and analysis techniques. Therefore, for better metaproteomics sample preparation, developing a new strategy or revising existing methods is necessary.

Microfluidics offer multiple advantages in the sample preparation of microorganisms for proteomic analysis, including automation, low-volume sample handling, safety and fast processing. Microfluidic technology has been successfully applied for the separation of bacterial and viral particles from bioaerosols (Hong et al., 2015), physical cell lysis (Grigorov et al., 2021 for a review) and chemical sample processing by utilizing immobilized trypsin (Huang et al., 2006). In a previous work (Ndiaye et al., 2020), we proposed a ChipFilter Proteolysis (CFP) microfluidic device as a reactor for the miniaturization of protein sample processing and digestion steps. The CFP design is closely related to the experimental setup of filter-aided sample processing (FASP). The microchip has two reaction chambers of 0.6 μl volume separated by a protein filtration membrane made using regenerated cellulose to concentrate or retain large polypeptides while releasing small molecules less than 10 kilodalton (kDa). Yeast protein extract and whole human cell proteome have been successfully analysed using CFP.

This study aims to assess a CFP-based workflow for sample preparation for LC-MS/MS analysis in the context of microbiology and metaproteomics. The workflow described for CFP (Ndiaye et al., 2020) was modified to introduce microbial cells directly into the device to perform all steps necessary for sample preparation starting from microbial cell lysis to proteolysis. On a mix of three microorganisms, CFP offers performance advantages compared to other methods including mFASP, in-gel and in-solution proteolysis. More proteins and peptides are identified with CFP than compared protocols, even at low cell density. The nature of the generated peptides was studied to better understand the influence of the microfluidic system in tryptic digestion. Finally, the CFP was utilized to prepare a sample mixture of 17 complex microbial species, leading to the identification of more than 10 species-specific proteins from 14 of the species.

## EXPERIMENTAL PROCEDURES

### Materials

Octyl-β-D-glucopyranoside (ODG), protease inhibitor, dithiothreitol (DTT), ammonium bicarbonate (ABC) and iodoacetamide (IAM) were purchased from Sigma-Aldrich. Trifluoroacetic acid was purchased from Thermo Fisher Scientific and acetonitrile (ACN) from Fisher Scientific.

Trypsin Gold, Mass Spectrometry Grade was from Promega. Lysozyme (50 ng/ml) was from Thermo Fisher Scientific.

Luria-Bertani (LB) agar (Merck) plates were inoculated with *Saccharomyces cerevisiae*, *Escherichia coli* and *Bacillus subtilis* and incubated at 37°C overnight. Inoculum into LB broth was made and cells were cultivated until the optical density reached 1 at 600 nm. Before harvesting, cells were counted using a glass slide under a microscope. The cells were collected by centrifugation at 2400 g for 5 minutes at room temperature. Single wash with phosphate buffer saline was performed prior to pelleting and stored at −80 ᵒC until further use. The three cell types were considered in different cell densities for the experiments. Two different dilutions of the cells were taken as a mix of 1:1:1 cell number ratio, 10E6 of each cell type for Mix 1 and 10E2 of each cell type for Mix 2. For Mix2, 10,000-fold dilution of the mix 1 was performed with 50mM ABC.

A standard whole-cell mixture consisting of 21 representative strains from 17 species of the gut microbiota (Zymo Research, ref D6331) was divided into 10 aliquots and stored in the storage solution provided by the manufacturer at −80 °C. This standard contains 18 bacterial strains including five strains of *E. coli* (JM109, B-3008, B-2207, B-766 and B-1109), 2 fungal strains, and 1 archaeal strain in staggered abundances, theoretically ranking from 20.01% to 0.0009% considering the cell number.

### ChipFilter method

The design and fabrication methodology of the microfluidic device has been explained previously (Ndiaye et al., 2020). For the comprehensive sample preparation, cells suspended in ABC were directly loaded into the device in a total volume of 30 μl. Cells were introduced into the ChipFilter using a piston syringe (Agilent) and syringe pump (Harvard Apparatus) maintaining a flow rate of 0.01 ml/minute. For the sequential injection of lysis buffer 1 [(1% (w/v) ODG, protease inhibitor in 150 mM Tris-HCl pH = 8.8)], 20 mM DTT in lysis buffer 1, 50 mM IAM in lysis buffer 1 and 50 mM ABC buffer was achieved using a flow-EZ pressure module, flow controller, M-switch (Fluigent), and the software Microfluidic Automation Tool (Fluigent). The flow rate and volume were maintained in two stages at 2 μl/minute for 45 μl and 1 μl/minute for 30 μl with the upper-pressure limit at 900 mbar. Finally, proteolysis was performed by introducing 20 μl of trypsin (final concentration of 0.1 μg/μl in 50 mM ABC) at room temperature. A constant flow of 50 mM ABC was maintained for 150 minutes to ensure mixing of the proteins with trypsin. The resulting proteolytic peptides were directly transferred in the flowthrough to the sample loop of the LC. The elution volume was regulated to capacitate sample loop volume. Proteolytic peptides were finally concentrated in a trapping column (C_18_ Pepmap, 300 μm i.d. × 5 mm length, Thermo Fisher Scientific).

For the standard gut microbiota, 75 μl of the mixture corresponding to approximately 3.94 * 10E8 cells were thawed in ice. For cell lysis, lysis buffer 2 that has the same composition as lysis buffer 1 with supplemented lysozyme (final concentration of 0.5 mg/ml) was used. All the subsequent steps were done as described above for mixed cells.

### Modified Filter-aided Sample Preparation (mFASP)

This protocol was modified from the original FASP protocol (Wiśniewski et al., 2009) to compare the two technologies (FASP and ChipFilter) that use very similar design under identical chemical conditions. Accordingly, the samples were resuspended in lysis buffer 1 (sample/lysis buffer 1 ratio of 1:5 v/v) and incubated at 37 ᵒC for 15 minutes. The contents are then transferred to Microcon-10 kDa Centrifugal Filter Units (Merck) and centrifuged at 15,000 g for 30 minutes to remove the flowthrough. Reduction was done with 20 mM DTT at 37 ᵒC for 2 hours and then the tubes were centrifuged at 15,000 g for 30 minutes with the flowthrough removed. Alkylation was performed with 50 mM IAM for 2 hours in the dark at 37 °C and the tubes were centrifuged at 15,000 g for 30 minutes to remove the flowthrough. Two washes with 500 μl of 50 mM ABC buffer were performed to remove the reagents. Digestion was performed with 2 μg trypsin in 50 mM ABC buffer for overnight at 37 °C. Elution of peptides was done by centrifugation twice at 15,000 g for 20 minutes with 50 mM ABC to ensure maximum peptide recovery. Finally, the peptides were acidified with 0.1% (v/v) TFA. Desalting was performed with C_18_ Zip Tips (Merck) as per manufacturer guidelines. Peptides were dried and stored in −20 °C until injection to LC.

### In-gel trypsin proteolysis

The samples were resuspended in 1X Laemmli buffer with β-Mercaptoethanol (Sigma Aldrich) and heated 5 minutes at 95 ᵒC. 40 μl of lysate were loaded into 12% precast gels (Bio-Rad) in Tris-Glycine-SDS running buffer at a voltage of 80V for 15 minutes. A short SDS-PAGE migration was used to restrict the proteome to a short band before separation (Hartmann et al., 2014). The gel was stained with Coomassie brilliant blue R-250 and the protein band was sliced and washed with excess double distilled water under agitation (700 rpm/minute) at room temperature. The gel was dehydrated with 100% ACN and dried using a Speedvac system. Reduction was performed with 10 mM DTT in 50 mM ABC buffer at 56ᵒC with agitation for 30 minutes. The excess solution was removed and the gel was dehydrated again with 100% ACN. The alkylation was performed next with 55 mM IAA in a 50mM ABC buffer with agitation at 37ᵒC for 20 minutes in the dark. The excess of solution was removed and replaced by a 50 mM ABC buffer. Dehydration was performed with 50% and 100% ACN and completely dried in the Speedvac system. 200 μl of trypsin (12.5 ng/μl) was prepared with 50mM ABC buffer and rehydration of the gel was done for 45 minutes at 4 ᵒC. Next, the excess solution was removed and digestion was performed for overnight at 37 ᵒC with agitation. Peptides were extracted from the gel in two steps by acidification and dehydration of the gel, using 100 μl of 1% TFA (v/v) and then 5% of formic acid (v/v) separately for 15 minutes under agitation at 37 ᵒC. The combined solutions were then transferred to a new tube. Again, the gel was dehydrated with 100% ACN for 15 minutes under agitation at 37 ᵒC and the solution was transferred to the elution tube. The eluate was dried in the Speedvac, resuspended in 20 μl of 1% (v/v) formic acid and sonicated for 5 minutes at 37 ᵒC. Desalting was performed with C_18_ zip tips as per manufacturer guidelines. Peptides were then dried and stored at −20 °C until injection to LC.

### In-solution Proteolysis

The samples were resuspended in Lysis buffer 3 (6 M Urea, 150 mM Tris, 1x Protease Inhibitors, 1% (w/v) octyl-β-D-glucopyranoside) with sample/lysis buffer 3 ratio of 1:5 (v/v). Samples were subjected to mechanical disruption with glass beads of 425-600 μm (Sigma-Aldrich) in the ratio of 1:1 (v/v). Five cycles consisting of 30 sec vortex and 60 sec in ice were repeated as bead-beating pre-treatment. The lysate was carefully removed from the beads and transferred to a new tube, where the reduction was performed with 20 mM DTT at 37 ᵒC for 2 hours. In the same tube, alkylation was performed with 100 mM IAA at 37 ᵒC for 2 hours in the dark. Precipitation of the proteins was performed with TCA solution (final concentration 10% v/v) for 30 minutes at 4 ᵒC, followed by centrifugation at 17,500 g at 4 ᵒC for 1 hour. The protein pellet was washed with 80% and then 100% (v/v) ice-cold acetone, followed by drying using speed-vac. The protein pellet was solubilized in a 50 mM ABC buffer and digestion was performed with 2 μg of trypsin for overnight at 37 °C. Acidification of the peptides was performed with 0.1% (v/v) TFA. Desalting was performed with C_18_ zip tips as per manufacturer guidelines. Peptides were dried after desalting and stored at −20 °C until injection to LC-MS.

### Liquid Chromatography and Mass Spectrometry

Samples were analysed by nanoLC-MS/MS in data-dependent acquisition (DDA) high-energy c-trap dissociation (HCD) mode using an RSLCnano UltiMate™ 3000 System coupled to a nanoESI Q-Exactive or Q-Exactive HF mass spectrometer (Thermo Fisher Scientific).

For the ChipFilter method, the peptides recovered in the trap column were separated on a capillary reverse-phase C18 column Pepmap 75 μm i.d. × 50 cm length (ThermoFisher Scientific) at 45°C with a linear 120 minutes gradient elution from 2.5% to 60% of buffer B (water/acetonitrile/formic acid 10%: 90%: 0.1% (v/v/v)) in buffer A (water/acetonitrile/formic acid 98%: 2%: 0.1% (v/v/v)) at a fixed flow rate of 220 nL/min.

For mFASP, in-gel and in-solution methods, the dried peptides were resuspended in 7 μl of 0.1 % TFA solution (v/v) and 6 μl was used for single-shot injection. Trapping was done with C_18_ Pepmap 300 μm i.d. × 5 mm length column (Thermo Fisher Scientific) and analysed in nanoLC MS/MS with a 120 minutes gradient as described earlier.

Mix-1 and Mix-2 samples analysis were performed with a Q-Exactive mass spectrometer operated in nanoESI at 1.7 kV. Full MS survey scan were recorded over the m/z range of 400 – 2000 with a resolution of 70,000 using an automatic gain control target value (AGC) of 3E6 with a maximum injection time of 100 ms. Up to 15 intense 2^+^ −5^+^ charged ions were selected for HCD with a normalized collision energy of 30, with a precursor isolation window at 2 m/z, resolution set at 17,500 with AGC value at 1E5 with a maximum injection time of 120 milliseconds. The minimum MS^2^ target value was set at 1E3 and dynamic exclusion for 20 seconds.

The standard gut microbiota was analysed with a Q-Exactive HF mass spectrometer operated in nanoESI (1.6 kV). Full MS survey scans were recorded over the m/z range of 375-1500 with a resolution of 60,000 using an automatic gain control target value (AGC) of 3E6 with a maximum injection time of 60 milliseconds. Up to 20 intense 2^+^ −5^+^ charged ions were selected for HCD with a normalized collision energy of 28%, with precursor isolation window at 2 m/z, resolution of 15,000, AGC value of 1E5 with a maximum injection time of 60 milliseconds. The minimum MS^2^ target value was set at 1E3 and dynamic exclusion for 20 seconds.

### Data Analysis

Spectra were processed using Proteome Discoverer (PD) v2.4 (Thermo Fisher Scientific). For mixed cells, the Mascot search engine (Matrix Science Mascot 2.2.04) was used against the all taxonomy SwissProt database (release 2022_03: 568002 sequences; 205171419 residues). For the standard gut microbiota, the SequestTM search engine was used against a dedicated sequence database (20,303 sequences containing 11,357,410 residues) specifically restricted to the species in standard gut mix from UniProtKB. A list 17 species from the standard gut mix includes - *Faecalibacterium prausnitzii*, *Roseburia hominis, Bifidobacterium adolescentis*, *Lactobacillus fermentum*, *Clostridioides difficile, Methanobrevibacter smithii*, *Enterococcus faecalis*, *Clostridium perfringens*, *Veillonella rogosae*, *Bacteroides fragilis*, *Prevotella corporis*, *Fusobacterium nucleatum*, *Akkermansia muciniphila*, *Salmonella enterica*, *Escherichia coli*, *Candida albicans* and *Saccharomyces cerevisiae* along with *homo sapiens* (1,133,353 sequences containing 506,763,197 residues). The database search was performed with the following parameters: MS and MS/MS mass tolerance 10 ppm and 0.02 Da respectively, trypsin specificity with up to 2 missed cleavages, partial Carbamidomethylation (C), Deamidation (NQ) and Oxidation (M). Proteins with at least one high confidence peptide and six amino acids were validated. Target FDR was set at 0.01. In order to perform label free quantification in PD, Minora feature detection tool was used in processing workflow and precursor ions quantifier in consensus workflow with consideration for both unique and razor peptides. The abundance value obtained was used to make the scatter plot and determine the correlation using tools in R (v 4.0.3).

The Kyte and Doolittle hydrophobicity index was calculated from a published package according to (Osorio et al., 2015).

All the experiments were performed in four replicates for mixed cells and in three replicates for standard gut mix. Statistical tests to identify significance (unpaired t test) were performed and values shown always represent mean ± standard error of mean. The mass spectrometry data have been deposed on the ProteomeXchange Consortium via the PRIDE partner repository (Perez Riverol et al., 2022) [px-submission #637484]

## RESULTS

### Cell lysis can be performed in ChipFilter without pre-treatment

In the previous work (N’Diaye et al., 2020), we have shown that it is possible to perform proteolysis of eukaryotic cell samples in the ChipFilter. In order to study microorganisms, it would be advantageous to perform the cell lysis directly in the ChipFilter too. It will help to avoid contamination issues and loss of material. To determine whether it was possible to lyse microbial cells on the chip, 1E6 cells of *E. coli, S. cerevisiae* and *B. subtills* were introduced separately in to the ChipFilter. Lysis was achieved using a lysis buffer containing 1% (w/v) ODG. After CFP, peptides were eluted and identified by mass spectrometry. The results are shown in Figure 1. The number of identified proteins ranked from 824 ± 50 (*B. subtills*), 1 121 ± 58 (*E. Coli*), to 1339 ± 44 (*S. cerevisiae*) suggesting that the ChipFilter cell lysis and proteolysis were efficient.

**Figure 1.**
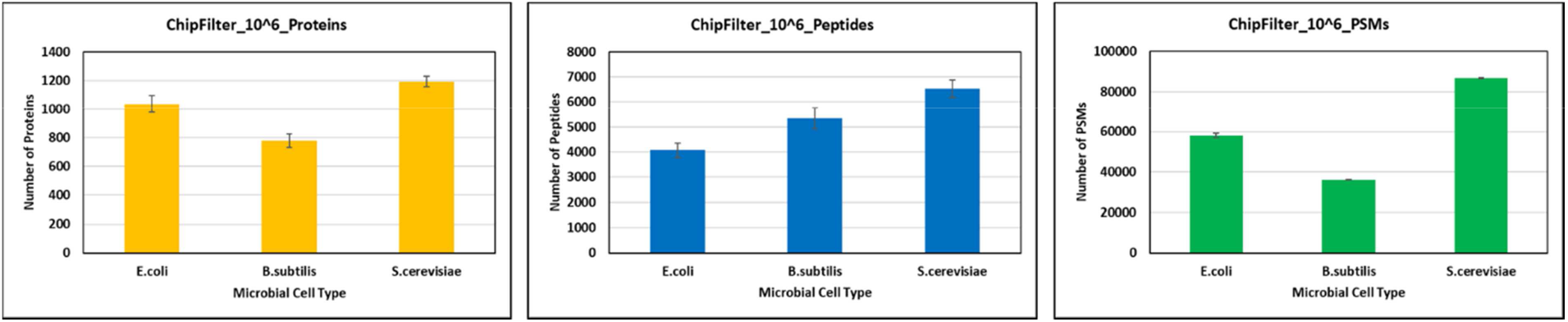
Whole cell lysis and bottom-up proteomic sample preparation with ChipFilter. Number of proteins (yellow), peptides (blue) and PSMs (green) identified for *Escherichia coli, Bacillus subtilis* and *Saccharomyces cerevisiae* using ChipFilter.

### ChipFilter method identifies more proteins and peptides than other methods

In order to assess the efficiency of ChipFilter for the identification of proteins from different microorganisms, a mixture of 1E6 *S. cerevisiae, E. coli* and *B. subtilis* (Mix 1) was analysed with CFP and standard proteolysis protocols (Figure 2A). The ChipFilter is a miniature of the FASP protocol, for which the fluid transfer is controlled by the laminar flow in the channels under an applied pressure, it is important to note that FASP tubes are designed for accommodating SDS based lysis buffers, but in the present study we used ODG based lysis buffers for FASP, called modified FASP (mFASP). This buffer and the lack of pre-treatment were chosen to keep it similar to the CFP. Next, an in-gel workflow is considered, as it is the most used workflow for metaproteomic sample preparation that allows the usage of SDS based lysis buffers. Finally, the in-solution method which introduces a cell lysis pre-treatment by bead beating was included to understand the influence of pre-treatment.

**Figure 2:**
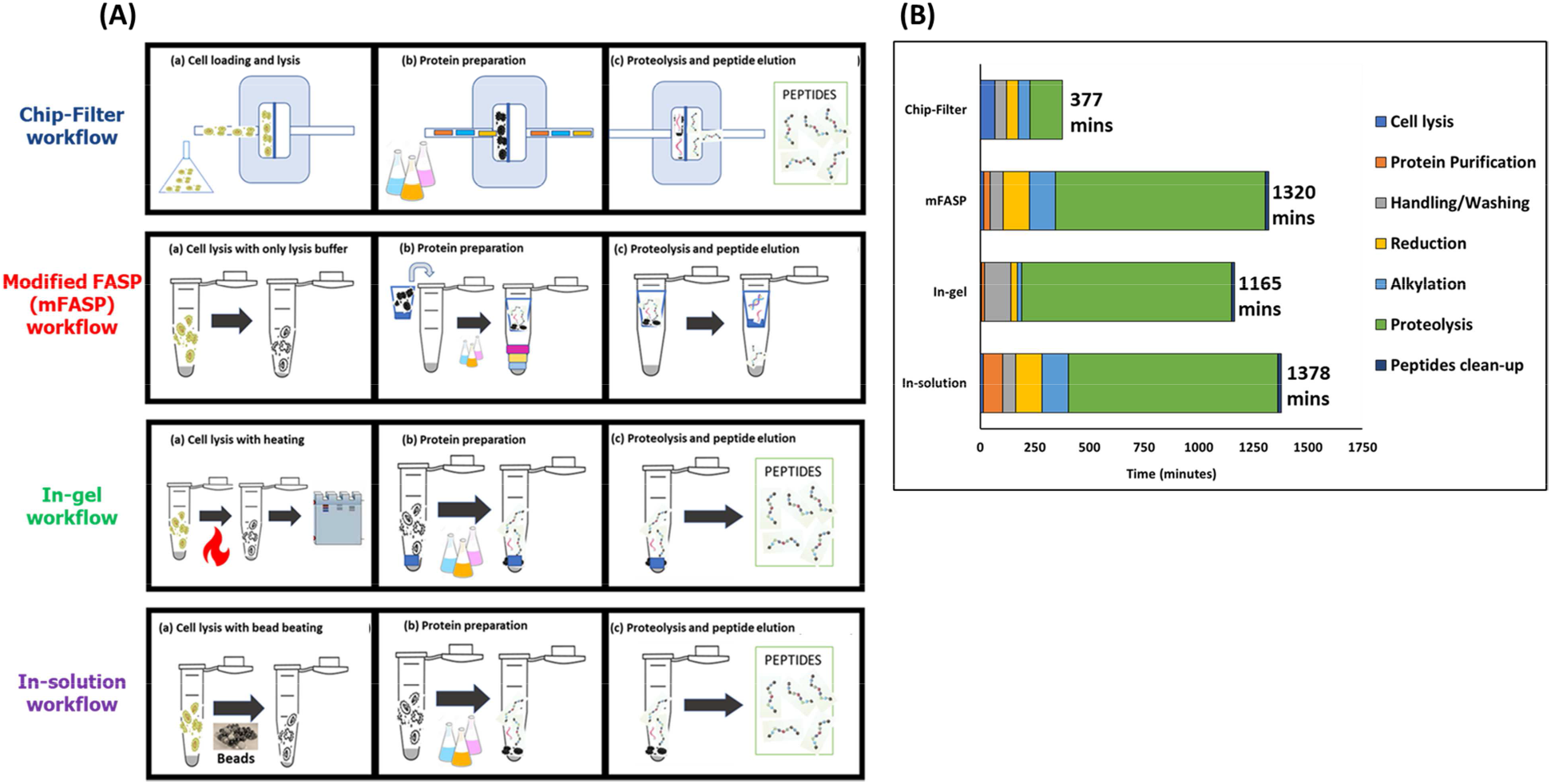
Comparative analysis of different workflows. (A) The different methodologies used to compare the performance of ChipFilter workflow includes modified filter assisted sample preparation (mFASP) workflow; protein separation by 12% SDS-PAGE gel and in-gel tryptic proteolysis; protein precipitation by TCA acidification and in-solution tryptic proteolysis. The steps involved in bottom-up proteomics are grouped into three key stages-cell lysis with or without pre-treatment; protein preparation including separation from other biomolecules and protein denaturation; trypsin mediated proteolysis. (B) The seven steps commonly used in bottom-up proteomics for each method along with the time taken for each step.

Figure 2 (B) highlights the time required for seven steps by each method considered. The seven steps considered include cell lysis, protein purification, handling step, reduction, alkylation, proteolysis and peptide clean-up. The digestion time for the ChipFilter is only two hours as compared to overnight for other methods, as used in most protocols. Shorter digestion time in the ChipFilter method is necessary to avoid the loss of peptides over time that exit the chamber upon digestion.

The performance of the protocols was evaluated by considering the number of target proteins and total peptides identified. For the concentration of 1E6 cells, the ChipFilter method achieved a mean protein identification of 1999 ± 54 whereas mFASP reached 1818 ± 91, in-gel 1504 ± 22 and in solution 1665 ± 15 (figure 3A). Similarly, at the peptide level, a significantly higher identification number is obtained for the ChipFilter method than other methods as shown in figure 3B. 9770 ± 108 peptides were identified using ChipFilter method, 8197 ± 435 with mFASP, 8218 ± 262 with in-gel and 7805 ± 178 with in-solution. Efficient catalysis in confined space and reduced loss of materials as compared to other methods can explain the higher identification for the ChipFilter method.

**Figure 3:**
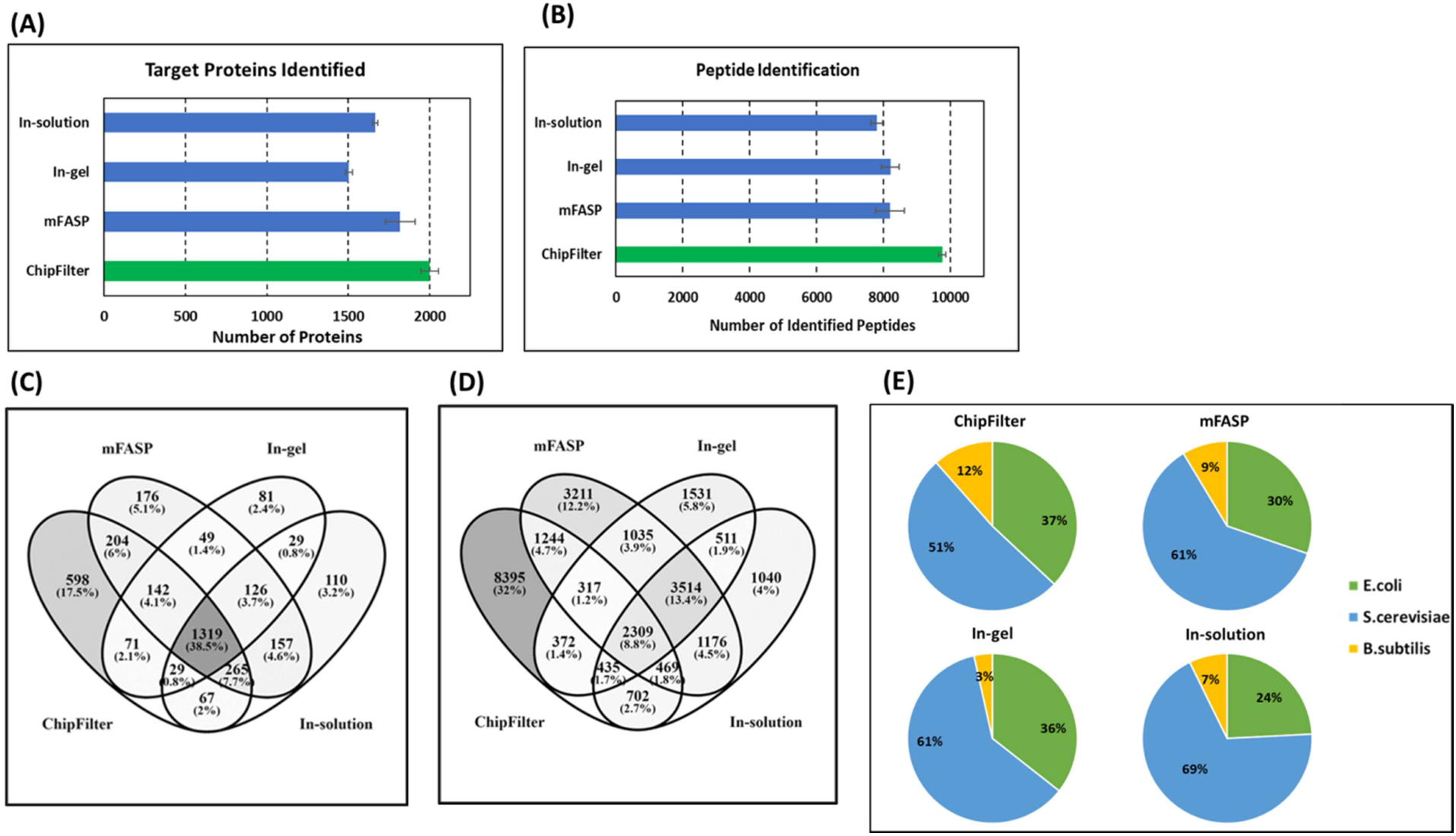
Performance assessment of ChipFilter workflow compared to other methods for Mix 1 (~3E6 cells/sample). Identification of (A) target proteins and (B) total peptides by different methods for four replicates. Distribution of (C) proteins and (D) peptides identified across different methods. (E) Percentage of proteins identified by each method for *E. coli*, *S. cerevisiae* and *B. subtilis*.

The distribution of the proteins and peptides among the methods show that ChipFilter protocol identifies the most distinctive proteins and peptides. At the protein level (figure 3C), 17.5 % (598 proteins) of the total proteins were identified exclusively by the ChipFilter method, whereas the other methods had less than 5.1% of specific proteins. The protein identification common between all the methods was about 38.5% (1319 proteins). At the peptide level (figure 3D), 32% (8395 peptides) of the total peptides were identified only in ChipFilter method. Interestingly, the peptide common to all the methods is only 8.8% (2309 peptides) of the total population, indicating that each method characteristically generate different peptide fractions. The ChipFilter allows the identification of 54.3% of the total peptides.

Comparison of the protein origin from three microbial cells (figure 3E) indicates that the ChipFilter identifies the highest percentage for gram-positive bacteria *B. subtilis* (12%) and gram-negative bacteria *E. coli* (37%) in comparison to other methods, whereas the highest percentage of fungi *S. cerevisiae* was recorded for in-solution (69%) method. Increased fungal proteins in the in-solution method can be reasoned due to the introduction of a pre-treatment step which was lacking in other methods. The in-gel method has poor identification for gram-positive bacteria. The usage of SDS containing Laemmli buffer can be the reason as the peptidoglycan cell wall of gram-positive bacteria was not efficiently denatured by SDS (Mahalanabis et al., 2009). The ChipFilter method can identify all the three cell types considerably better than other methods without pre-treatment.

### The ChipFilter method performs efficiently with low cell number

ChipFilter was successfully used on a sample with low numbers (3E2) in mix 2 of the three microorganisms. The number of identified proteins was similar to the number obtained after in-solution proteolysis, and significantly higher than results obtained with in-gel and mFASP protocols (Figure 4A). At the peptide level, ChipFilter method allowed the identification of a greater number of peptides than other three methods tested (Figure 4B).

**Figure 4:**
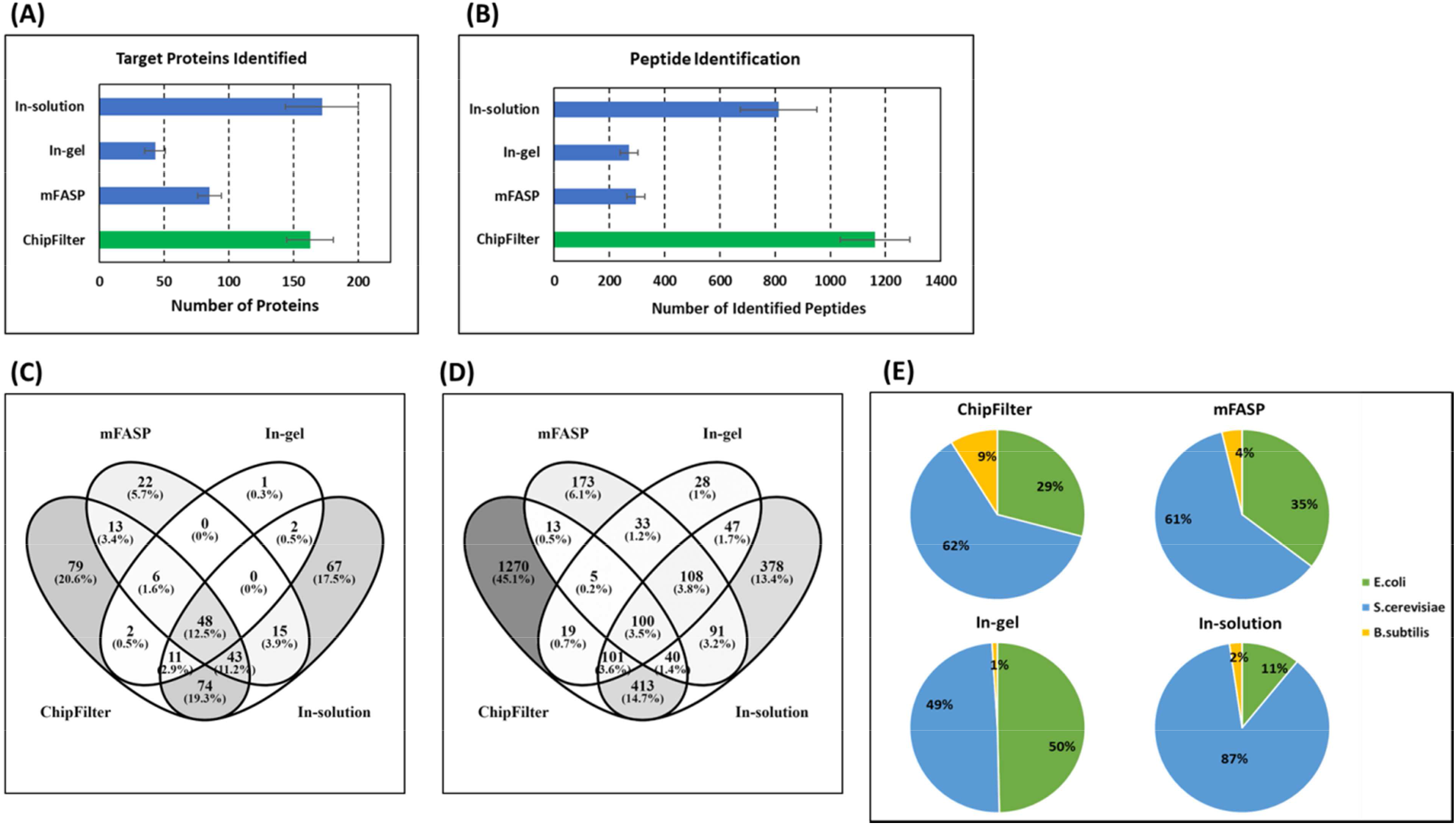
Performance assessment of ChipFilter workflow compared to other methods for Mix 2 (~3E2 cells/type). Identification of (A) target proteins and (B) total peptides by different methods for four replicates. Distribution of (C) proteins and (D) peptides identified across different methods. (E) Percentage of proteins identified by each method for *E. coli*, *S. cerevisiae* and *B. subtilis*.

The sensitivity of ChipFilter method was tested for low cell number sample (Mix 2) to assess the identification and distribution of peptides and proteins like in mix 1. The mean number of target proteins and total peptides identified are 163 ± 18 and 1162 ± 126 respectively (figure 4A and 4B). The number of protein identifications is similar for ChipFilter method as compared to in-solution method (172 ± 28 proteins), but the number of peptide identification is higher than every other method.

As observed at 3E6 cell density, the coverage of identified proteins is variable depending on the protocol used. For Mix 2, the ChipFilter method identified the highest proportion of proteins (72% of the total) and the highest proportion of exclusive proteins (20.6% of the total target protein identification; Figure 4C). The same trends are obtained at the peptide level. The ChipFilter method identifies the majority of peptides (69.7%), and is the protocol that allows the identification of the largest number of exclusive peptides (45.1% of the total, Figure 4D).

The distribution of the identified proteins according the species is more balanced with ChipFilter protocol, in particular for *Bacillus* proteins which represent 9% of the total (9 proteins) against 2% for the other protocols (less than 4 proteins, Figure 4E)

The distribution of the proteins (figure 4C) indicates that the percentage of proteins exclusively identified by ChipFilter method (20.6%, 79 proteins) and in-solution method (17.5%, 67 proteins) are very close, unlike in mix 1. Furthermore, the percentage of proteins identified by both methods is about 45.9%, suggesting that the protein representatively is very similar. Of the total proteins identified, only 27.9% were not identified by the ChipFilter method. For the peptide identifications shown in figure 4D, the peptides exclusively identified by the ChipFilter method is 45.1% (1270 peptides). It is roughly three times higher than the peptides exclusively identified by in-solution method (13.4%, 378 peptides). The proportion of peptides not identified by the ChipFilter method is 30.4% (858 peptides). As observed with Mix 1, the distribution of peptides or proteins is not similar, indicating that sample preparation methods handle low cell density sample differently. Finally, the representation of protein identifications for different species with the methods is indicated in figure 4E. Like in Mix 1, the distribution of the proteins identified is represented for all the three species for the ChipFilter method. In the case of identification of *B. subtilis*, the ChipFilter identifies 9% of total proteins whereas, it is less than 4% for the other methods. Furthermore, the protein identification for *S. cerevisiae* for the in-solution method is 87% of the total proteins identified which is higher than the other methods which have about 50% to 60% of this species. The ChipFilter method clearly is a universal sample preparation method for simultaneously identifying proteins in a single analysis even at low cell numbers.

### Repeatability in ChipFilter workflow is high

To assess the repeatability of ChipFilter method, LFQ intensities among the four replicates of Mix 1 and Mix 2 cell samples have been studied. The correlation coefficient values are in the range of 0.76 to 0.83 for Mix 1 (Figure 5A), and in the range of 0.59 to 0.69 for Mix 2 (Figure 5B). The lower correlation can be due to the decreased intensities and signal-to-noise obtained for low cell numbers. These correlation coefficients are comparable to those obtained for other methods with slightly lower values for ChipFilter method.

**Figure 5:**
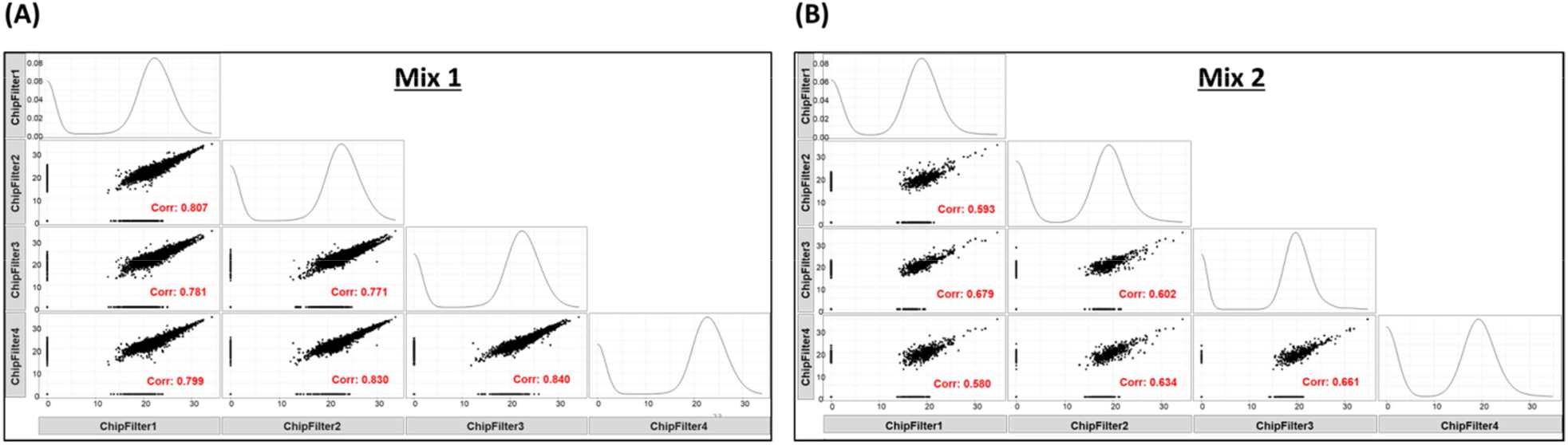
Association between the replicates for ChipFilter method for mix 1 and 2. The LFQ intensity is used to show the association between the different replicates (A) for mix 1 and (B) mix 2 (B). The Pearson’s correlation value is indicated in red for every pair along with the scatter plot distribution.

Repeatability is often difficult to achieve in metaproteomics samples but is critical. For the current study, the variations arising during nanoLC-MS/MS are not considered and are attributed to the sample preparation methodology. Correlation is shown to be in the positive direction, indicating that changing the parameters in one replicate induces similar changes in other replicates. The correlation coefficient values for the ChipFilter method indicate a high positive correlation between the replicates for high cell numbers. Even though the peptide identifications are significantly high for the ChipFilter method in Mix 2, the correlation between the replicates make this method useable. The reduction in the correlation between the two cell densities can be due to the reduced intensities obtained at lower cell number. Comparing the correlation coefficients between the methods (data not shown) indicates that the ChipFilter method has high variability among the replicates. The in-solution and in-gel methods exhibit lower standard deviation than other methods. mFASP generates high variability like the ChipFilter method and it can be possible that the filter-based methods could contribute to these differences. In conclusion, the ChipFilter method generates replicates that have positive correlation in Mix 1 and 2 indicating that reproducibility between the different cell numbers is related to each other in the same manner.

### Physio-chemical characteristics of the peptides generated by the ChipFilter method

As the ChipFilter method allows the identification of a large number of specific peptides (Figure 2D and 3D), these peptides were studied to see if they have different physio-chemical characteristics from the peptides obtained by other methods, which could lead to a bias in the exploitation of the results. This comparison was done for 3E6 and 3E2 cell samples.

For the ChipFilter method with Mix 1 sample, 32% (3137 ± 149) of the peptides have a mass below 2000 Da, whereas for other methods it is approximately 73% (5879 ± 232 peptides) (Figure 6A). For Mix 2 sample, the percentage of peptides with a mass below 2000 Da was 56% in the ChipFilter. It is to be noted that the highest mass of the peptides generated was 4988 Da for a ChipFilter cut-off of 10,000 Da. These results are in line with those for amino acid size (Figure 6B). Using the ChipFilter method, 42% of Mix 1 peptides have a length of up to 20 amino acids, instead of approximately 80% for other methods. Similarly, the proportion of Mix 2 peptides shorter than 20 amino acids was 66% with the CFP while 91% for the other methods.

**Figure 6.**
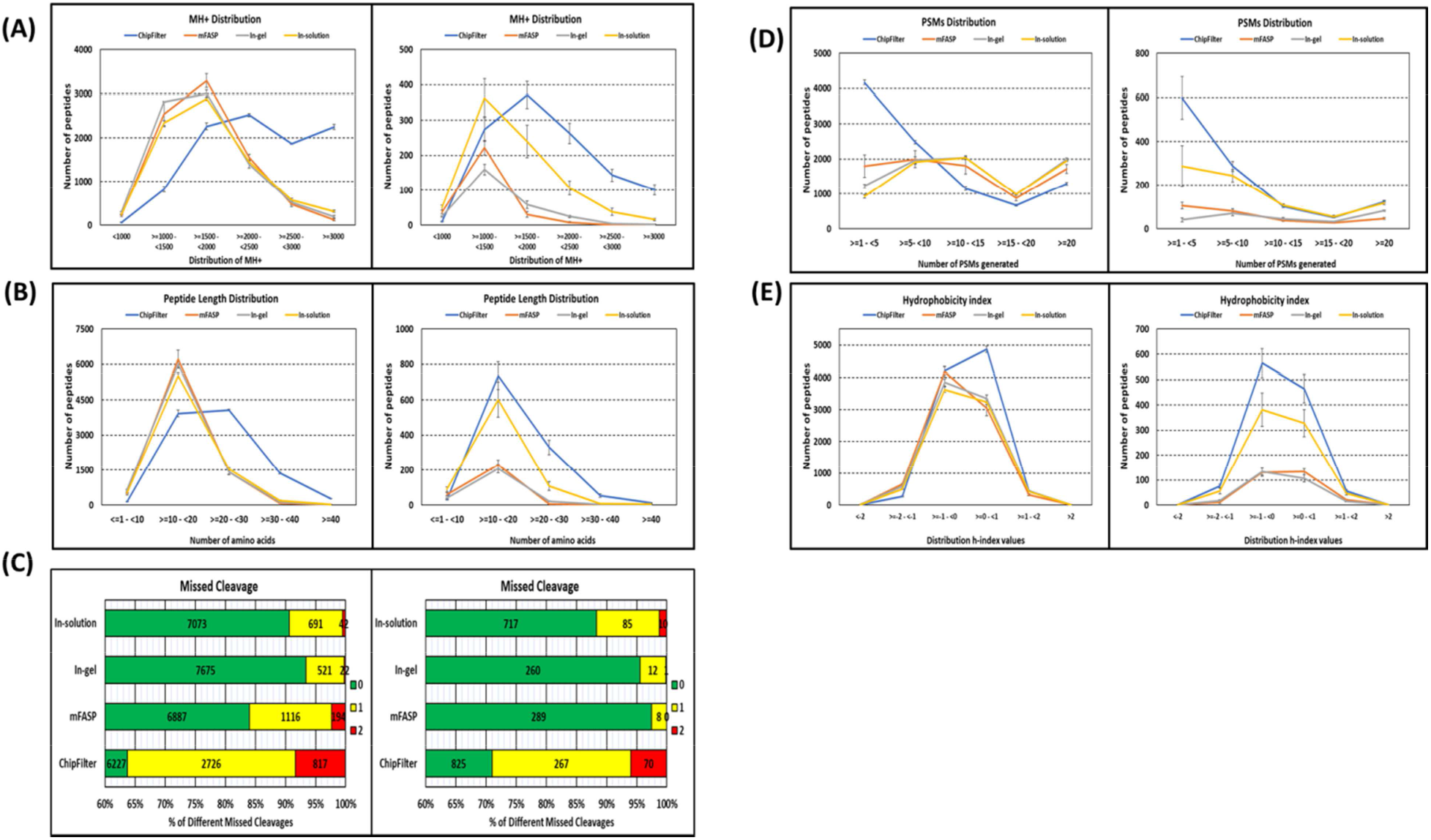
Physical chemical characteristics of the peptides generated by different preparation methods. (A) Positive ion mass (MH+) distribution for Mix 1 (left) and Mix 2 cell (right) samples. (B) Amino acid length distribution for Mix 1 (left) and Mix 2 (right) cell samples. (C) Ratio of missed cleavages generated by each method for Mix 1 (left) and Mix 2 (right) cell samples. (D) Peptide spectral matches (PSMs) distribution for Mix 1 (left) and Mix 2 (right) cell samples. (E) Kyte-Doolittle hydrophobicity index is shown in (E) for Mix 1 (left) and Mix 2 (right) samples. For (B) to (E) the mean value is plotted in the graph along with the error bars indicating the standard deviation between four replicates.

To explain this result, the percentage of missed cleavage was studied for Mix 1 and Mix 2 samples (Figure 6C). The ChipFilter method has the highest percentages of 1 and 2 missed cleavages with Mix 1 (28% and 8% respectively) and Mix 2 (23% and 6% respectively). The percentage is also considerably high in the mFASP method for Mix 1 (14% and 2% respectively). In-solution method is consistent between the two cell numbers with around 10% for 1 missed cleavage and 1% for 2 missed cleavages. The in-gel method has the least. The higher proportion of missed cleavage with ChipFilter can be explained by factors such as higher density of proteins in a confined volume or shorter proteolysis duration.

Peptide spectral matches (PSMs) are useful to identify the sequence of the peptide’s spectra generated in tandem MS by assigning scores to the theoretical spectra that can be generated for a specific peptide fragment. This method depends on the fragmentation of the peptides to compute the scores. Often it is considered that generation of a high number of PSMs per peptide is appropriate. On this note, the PSMs distribution was studied and is represented in figure 6D. In Mix 1, results indicated that the fraction of peptides identified with less than 10 PSMs is 6649 ± 112 (68%) for the ChipFilter method, whereas in other methods, this number dropped to 3273 ± 272 (36%). In Mix 2, the fraction of peptides under 10 PSMs is higher for the ChipFilter method (844 ± 119 peptides, 76%) than in Mix 1. Interestingly in Mix 2 except for in-gel (41%), other methods generated higher fraction of peptides under 10 PSMs (68%) The higher percentage of missed cleavages in the ChipFilter method could have contributed to the generation of peptides with a lower PSM number.

Finally, the hydrophobicity of the peptides generated was studied using the Kyte-Doolittle hydrophobicity index (figure 6E). Most of the peptides were accumulated between −1 to +1 which is indicative of a neutral mixture with slight hydrophobic or slight hydrophilic peptides. This trend was observed for all the methods including the ChipFilter method for both cell mixtures.

### ChipFilter allows efficient proteomic analysis of a standard gut microbiome

In order to assess the efficiency of ChipFilter proteolysis on a more complex sample, a commercial mixture of 17 species, including bacteria (Gram-negative and Gram-positive), fungi and archaea, was used. This gut standard mixture is non-uniform in cell number and contains species commonly found in the intestinal microbiota of humans.

A mean of 2521 ± 375 proteins belonging to the 17 expected species have been identified (Figure 7) with a total of 6099 ± 1140 peptides and 18 339 ± 1535 PSMs.

**Figure 7:**
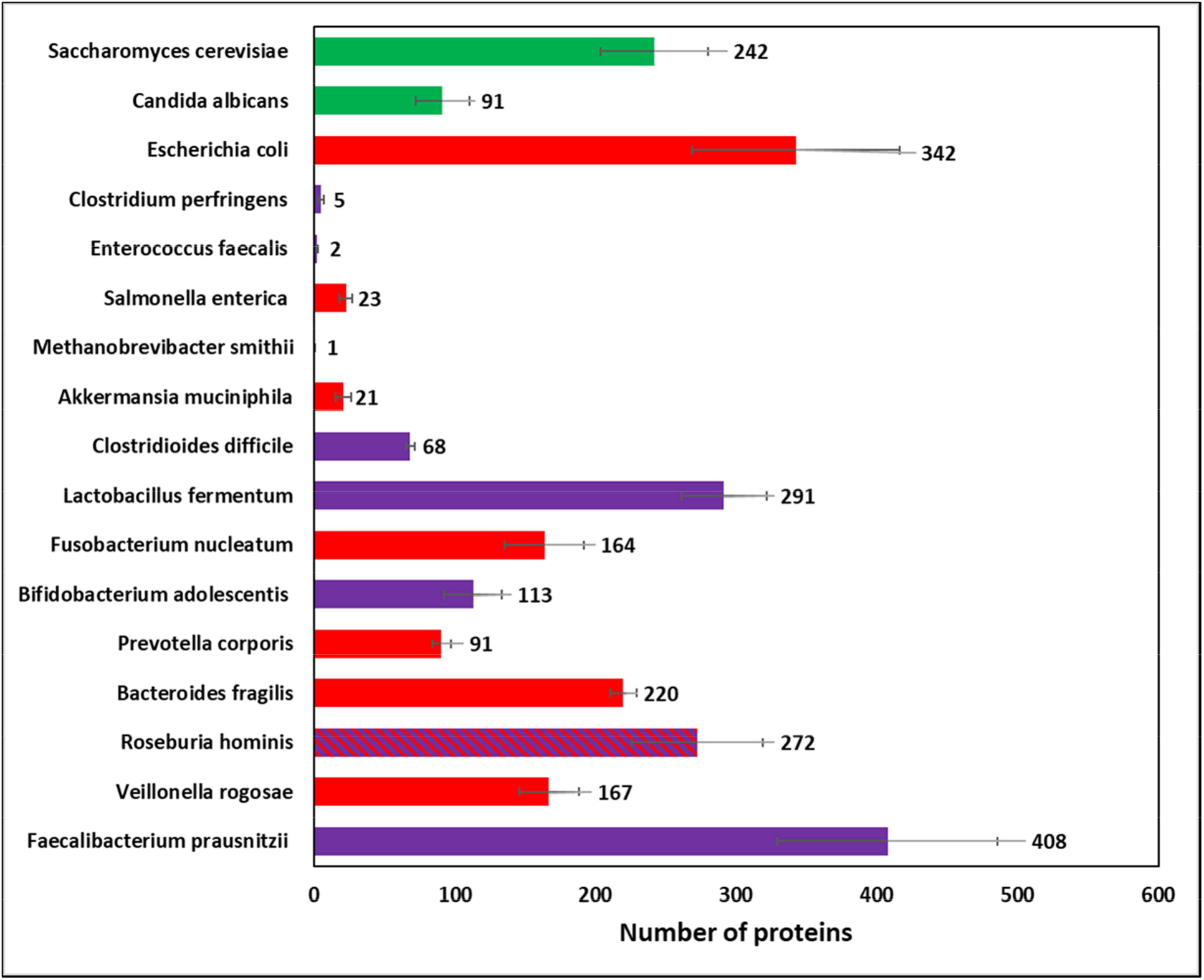
Number of proteins identified from each species in the standard gut mix after preparation by CFP. Green: fungi, Red: Gram-negative bacteria; Violet: Gram-positive bacteria.

*R. hominis* which is Gram invariable is represented in two colours. The five separate *E. coli* strains present in the mixture are grouped under a single species name.

The proteomic data were used to estimate the biomass of each species (Kleiner et al., 2017). The results are compared with the cell number, 16S and 18S and genome copy number information provided by the manufacturer (Figure 8).

**Figure 8:**
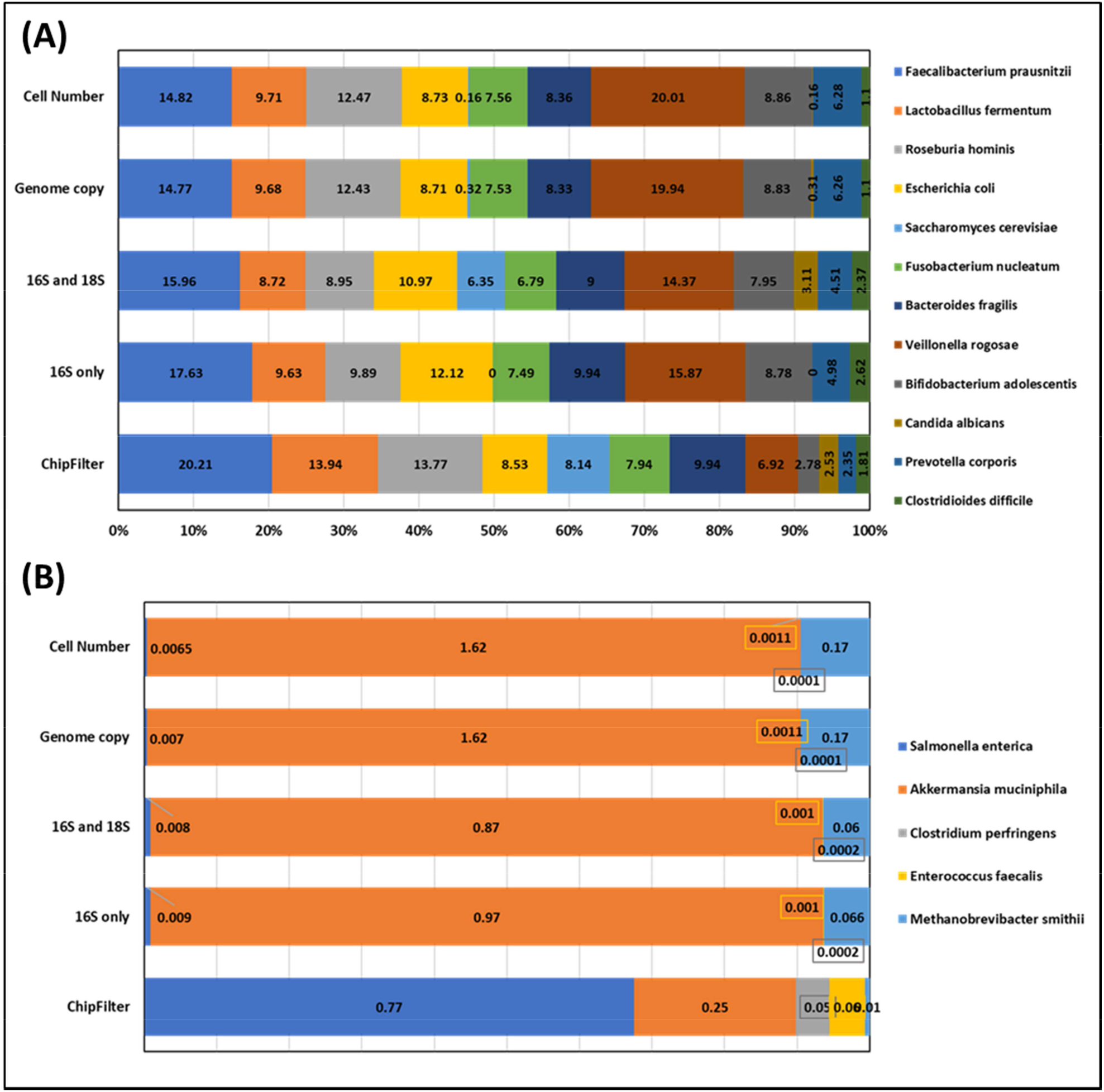
Biomass estimations using Next Generation Sequencing methods and ChipFilter proteomics analysis, for species present in the standard gut mix at (A) at more than 1% abundance and (B) less than 1% abundance (B). *Cell number, genome copy, 16S and 18S and 16S data were provided by the manufacturer.

## DISCUSSION

In previous work, we have shown the usefulness of the ChipFilter device for the preparation of eukaryotic cells for bottom-up proteomic analysis (Ndiaye et al., 2020). In the present study, we have adapted this system to the analysis of microbial cells, having different phenotype and cell wall compositions. We have shown that it is possible to perform cell lysis directly into the ChipFilter device, before other pre-analytical steps are performed. On a mixture of three species, we identified more proteins and peptides than standard protocols compared in the study. We have extended the use of this ChipFilter device to the analysis of a standard gut microbiome. Proteins from almost all species were identified. Moreover, these results were be used to estimate the contribution of each species to the biomass, giving results in line with those determined by nucleic acid techniques.

To use the ChipFilter device for whole microbial cell sample preparation, we had to make two types of changes. The first concerns the sample loading. Our earlier work mainly focused on the usage of cellular lysates that was introduced to the ChipFilter by using the same microfluidic capillaries used for the other reagents. When using this set-up for microbial cells, we observed a cross-contamination between the samples. To overcome this problem, samples were introduced in the device using an external piston syringe and pump, and not through capillaries. Second, the lysis buffer containing ODG was adapted to be more efficient in microbial cell lysis with addition of lysozyme in case of complex mixtures comprising Gram positive bacteria. Detergents are the commonly preferred reagents for cell lysis which are often enhanced by use of physical or mechanical lysis methods to break the cell wall and membranes. Commonly used lysis buffers use anionic detergent-sodium dodecyl sulfate (SDS) in the range of 1 – 5% (m/v). SDS is highly effective in cell lysis but is generally associated with several drawbacks including requirement for urea washing to neutralize, reduction in trypsin activity and influence in mass spectrometry detection (Botelho et al., 2010). Hence, we use an alternative lysis buffer without SDS, that contains non-ionic detergent ODG. It allowed direct cell lysis on the microfluidic device without any pre-treatment. Simple washing with ammonium bicarbonate buffer was sufficient to remove the detergent from the device.

Particular attention was paid to the physio-chemical characterisation of the peptides produced after CFP. In CFP, variation in the nature of peptides generated can be caused by peptide adsorption on PDMS, continuous fluid flow and protein processing conditions provided. Peptides generated by the ChipFilter method were novel as they had higher percentage of peptides with long amino acid chains, higher mass and more missed cleavage than other methods included in the study. One of the reasons for the generation of distinct peptides can be the proteolysis condition provided inside the ChipFilter. Mechanisms of proteolysis can be broadly classified into in-gel (Shevchenko et al., 2006), in-solution (León et al., 2013) and in-solid phase (Hughes et al., 2019). While the mechanism of catalysis in the device can be argued to be a hybrid between in-solution and in-solid phase. The chamber offers the space for interaction between the proteins and the trypsin similar to in-solution, whereas the nitrocellulose membrane and the PDMS adsorb the trypsin thereby acting as solid-phase to enable catalysis. Also, during the catalysis, a small flow rate (0.5 μl/minute) of the ABC buffer was provided to increase the exchange of the reactants. These dual digestion mechanisms offered by the device can also be an influencing factor in the generation of distinct peptides. It is well known that shorter digestion time can generate missed cleavages. Further, shorter digestion times have been reported in past study to be advantageous (Deng et al., 2018). The ChipFilter also identifies most of the peptides identified by other methods. All this suggests that even though the number and size of peptides is greater after ChipFilter proteolysis, it does not seem to induce an identification bias compared to other methods.

The ChipFilter method was used to study standard gut microbiome consisting of 17 different species with different abundance. Our proteomic data allowed us to identify at least 10 proteins specific to 14 of these species. Proteins support the vast majority of the metabolic functions in living organisms. Knowing the protein composition of a community allows to distinguish the nature of the microorganisms that generate it and is necessary to describe functions like cell-to-cell interactions, contribution of cells in the biochemical pathways operating within a community like breakdown of food by gut communities (Wang et al., 2020) or identifying biomarkers (Heintz-Buschart et al., 2016; Park et al. 2021). In the present work, functional analysis of identified proteins has not been carried out due to use of standard mixture. But as a proof of concept, the contribution of each species in the overall biomass was quantified according to (Kleiner et al. 2017). In this model, the quantification data are summed based on the taxonomic assignment of inferred proteins and not based on the taxonomic assignment of peptide identifications because peptides are frequently associated with multiple proteins from different taxa. Our results are in line with those proposed by other sequencing (16S, 18S or shot-gun sequencing) results provided by the manufacturers. This aligns the proteomic finding with nucleic acid results obtained externally. It also shows the feasibility of the ChipFilter method for proteomic sample preparation.

In conclusion, the ChipFilter based sample preparation method for bottom-up proteomics by LC-MS/MS allowed in the identification of microbial proteins from single species to communities, thereby establishing the feasibility for metaproteomics. The advantage to confine low cell number cells without loss of proteins during lysis or washing stages, automation and efficient identification will allow in accelerated sample preparation with better identification. The use of ChipFilter could be extended to study microbiomes with low sample volume or cell density that will require confinement during preanalytical steps.

## ACKNOLEDGMENT

This project has received funding from the European Union’s Horizon 2020 research and innovation programme under the Marie Skłodowska-Curie grant agreement no. 754387, the PSL University and Région Ile de France.

This work has benefited from the technical contribution of the joint service unit CNRS UAR 3750. The authors would like to thank the engineers of this unit for their advice during the development of the experiments.

